# Reliable Evaluation and Learning in Multi-input Biological Association Prediction

**DOI:** 10.64898/2026.02.17.706287

**Authors:** Sobhan Ahmadian Moghadam, Hesam Montazeri

**Author notes:** Correspondence to Hesam Montazeri.

## Abstract

Multi-input association prediction is central to many key problems in computational biology, spanning tasks from drug-target and protein–protein interactions to higher-order challenges such as drug synergy modeling and MHC–peptide–TCR binding. Yet widely used benchmarks often overestimate performance by enabling models to exploit degree ratio shortcuts, while alternative out-of-distribution splits are overly restrictive and impractical. Here we introduce an entity-balanced evaluation framework that systematically neutralizes shortcut signals by balancing positive and negative associations at the entity level. This enables fairer assessments that reflect genuine relational learning and extend naturally from pairwise to multi-entity problems. We further present UnbiasNet, a model-agnostic training strategy that cycles through diverse entity-balanced sub-training sets, removing access to degree ratio bias and enhancing robustness. Applied to drug–target interaction and drug synergy prediction, our framework reveals the extent of shortcut reliance in existing methods while enabling consistent identification of meaningful biological associations, thereby setting a rigorous foundation for future methodological progress.

## Introduction

Rigorous evaluation frameworks have been central to progress in computational biology, as demonstrated by community-wide benchmarks such as the Critical Assessment of Protein Structure Prediction (CASP)^1^ or the Critical Assessment of protein Intrinsic Disorder prediction (CAID)^2^. Such benchmarks show how well-designed datasets and evaluation metrics can reveal the specific strengths and weaknesses of competing methods^2,3^. By contrast, many other computational biology tasks lack standardized unbiased assessments, leading to inflated performance claims and benchmarks misaligned with true generalization, which in turn impedes genuine methodological advancement.

A broad class of problems particularly affected by this limitation are *multi-input association prediction tasks*, where the goal is to determine whether an association exists between biological entities (Figs. 1A, 1B). In the simplest case, binary (X–Y) associations include tasks such as drug–target^4–6^, protein–protein^7–9^, miRNA-disease^10,11^, RNA-protein^12,13^ or phage-host^14,15^ interaction prediction, where X and Y denote two distinct types of biological entities (e.g., drugs and proteins). More complex, higher-order associations arise in settings such as drug synergy modeling^16^ or MHC–peptide–TCR^17^ binding prediction. The number of input entities can thus range from pairs to triplets or larger sets, but the output is a binary variable indicating the presence or absence of an association. To address these tasks, a wide variety of computational methods have been developed, spanning deep learning models such as MIDTI^18^ and MHGNN^19^ for drug–target interaction and CCSynergy^16^ and TranSynergy^20^ for drug synergy prediction.

**Fig. 1.**
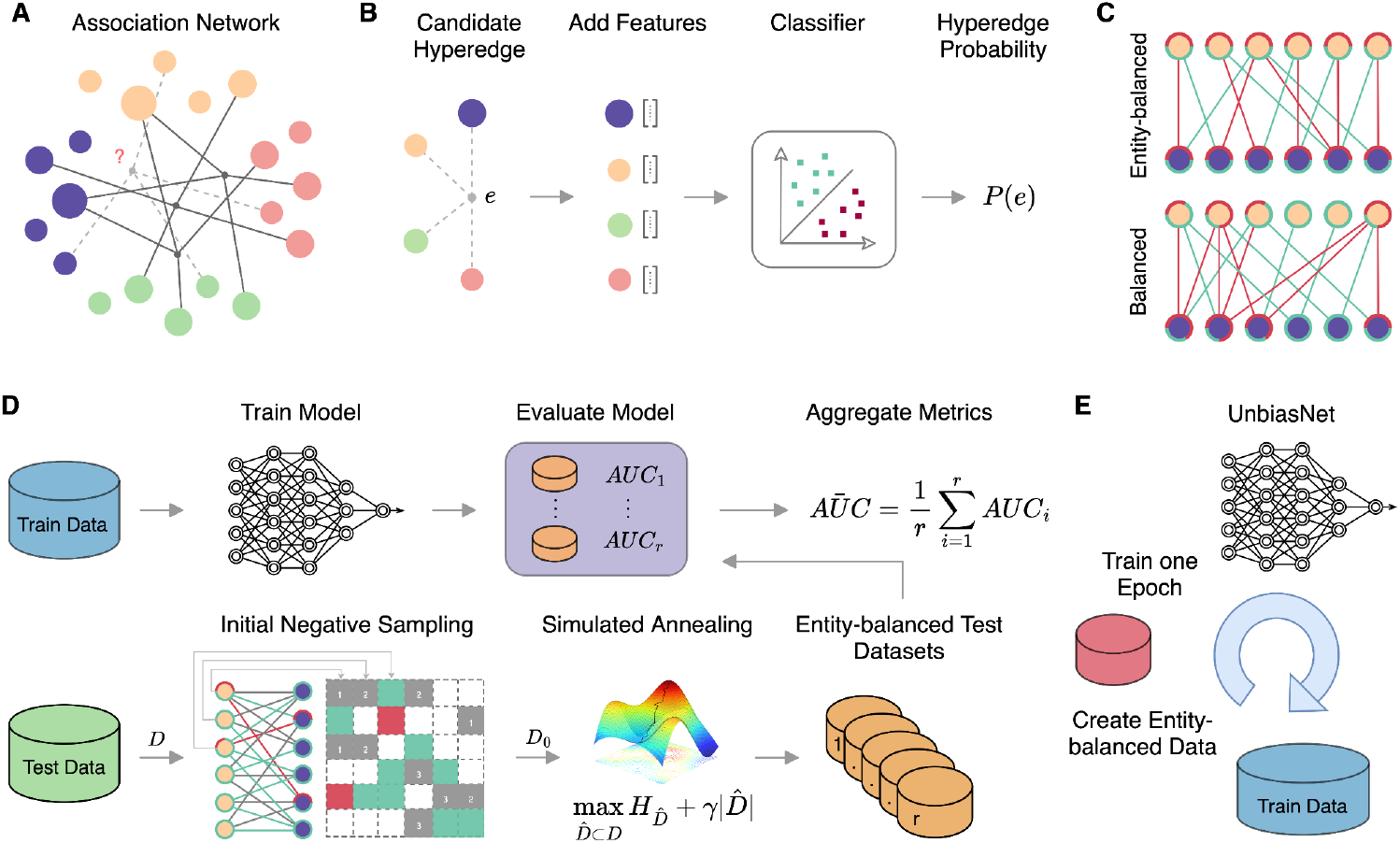
Overview of multi-input association prediction, evaluation challenges, and the proposed solution. **A**, Example of an association network with four entity types. **B**, In multi-input association prediction, the task is to determine whether a specific hyperedge (association) exists, based on numerical features extracted for each participating entity. **C**, Balanced vs. entity-balanced networks: both have equal positives and negatives, but only the latter achieves per-entity balance (the red/green indicators encircling the nodes). **D**, Entity-balanced evaluation framework. Models are trained as usual but evaluated on entity-balanced test sets generated through two steps: degree-guided negative sampling followed by simulated annealing using an entropy-based score. **E**, UnbiasNet: training across multiple entity-balanced datasets removes shortcut access and improves generalization.

Despite progress, conventional evaluation frameworks for association prediction tasks often mask a critical vulnerability: shortcut learning^21^. This occurs when models exploit spurious statistical regularities in the data rather than learning meaningful biological patterns. In the context of multi-input association prediction, the most pervasive shortcut is degree ratio bias—the reliance on the balance of known positive and negative associations linked to each entity^22–24^. While leveraging this superficial unintended feature can yield seemingly strong results on conventional benchmarks, such models fail to generalize to real-world scenarios. Moreover, designing models that avoid such shortcuts remains a central challenge. For example, Chatterjee et al.^22^ proposed a method for protein–ligand prediction, which samples negatives from distant nodes in the interaction graph to mitigate degree ratio reliance. Yet, the precise mechanism underlying its robustness is not well understood, and the approach remains restricted to binary associations.

Conventional evaluation frameworks, which typically rely on random train–test splits and metrics such as AUC, are especially susceptible to the degree ratio bias. Even when global class balance is enforced by subsampling negatives, the distribution of associations across individual entities remains highly polarized—some entities are linked almost exclusively to positives, others to negatives (Fig 1C). This entity-level imbalance, a form of Simpson’s paradox, preserves shortcut signals across training and test sets. As a result, models can exploit degree ratio instead of learning genuine relational features, yet still achieve deceptively high performance scores. As demonstrated in prior studies^22,23^ and reaffirmed by our results, naïve baseline models that rely solely on degree ratio can equal or even surpass the performance of state-of-the-art methods under conventional evaluation protocols, revealing shortcomings in particular evaluation practices that remain widely used in association prediction.

One strategy to mitigate this problem is to stratify the test dataset based on whether entities in test associations are also present in training data^22–25^. This creates an out-of-distribution (o.o.d.) test set, where the test entities never appear in training, preventing models from exploiting degree ratio information. However, this strategy quickly becomes impractical. For example, graph-based methods rely on the connectivity of the association network to learn meaningful patterns, making evaluation infeasible if the test set entirely excludes training entities. Extending this approach to higher-order associations (e.g., triplets) is overly restrictive, as ensuring disjoint entity sets across training, validation, and test splits severely reduces usable data. Moreover, models can estimate an entity’s degree ratio by leveraging other entities that are sequence-similar to the query^24^. Consequently, this type of evaluation also requires controlling for sequence similarity between the test and training data, which further complicates the problem and makes the approach overly conservative for realistic evaluation.

To tackle these problems, we propose a sampling algorithm for constructing entity-balanced out-of-distribution (o.o.d.) datasets (Figs. 1C, 1D). The process begins with iterative sampling guided by the degree ratio distribution, which increases the likelihood of selecting candidate negative edges that reduce entity-balance polarity. Next, we apply simulated annealing using an entropy-based score function to further mitigate polarity bias. The probabilistic nature of this procedure enables the generation of diverse entity-balanced datasets. We then assess model performance across multiple such datasets and aggregate the results (see Methods). As we will show, this framework discourages reliance on shortcuts, ensures fair evaluation even when entities overlap between training and test sets, removes the need for explicit similarity controls due to degree-ratio bias, and generalizes naturally to multi-input prediction tasks.

Once biases in the data are identified, a central challenge lies in designing models that remain robust to them^26^. To address this for the degree ratio bias, we introduce UnbiasNet, a learning framework explicitly developed to counteract shortcut learning in deep neural networks (Fig. 1E). To mitigate degree ratio bias, UnbiasNet employs multiple entity-balanced sub-training sets, presenting the model with a different dataset at each epoch. This strategy diversifies the learning signal, systematically removes access to degree ratio, and enhances generalization and robustness. Importantly, the framework is model-agnostic and can be applied across diverse deep learning architectures and association prediction tasks.

## Results

We focused on two canonical association prediction problems: drug–target interaction (DTI) prediction and drug synergy modeling (Figs. 2-3). For each task, we benchmarked one state-of-the-art model alongside conventional classifiers (Random Forest, XGBoost, Linear Regression and MLP) trained on the same feature representations. Evaluations were conducted under three distinct frameworks: (1) Full Test Evaluation, reflecting the natural class imbalance; (2) Balanced Evaluation, which equalizes positive and negative samples; and (3) our proposed Entity-Balanced Evaluation, which explicitly controls for degree ratio shortcut learning. All evaluations were performed using repeated cross-validation, with differences arising only from the strategy used to sample and balance the test sets (see Methods).

**Fig. 2.**
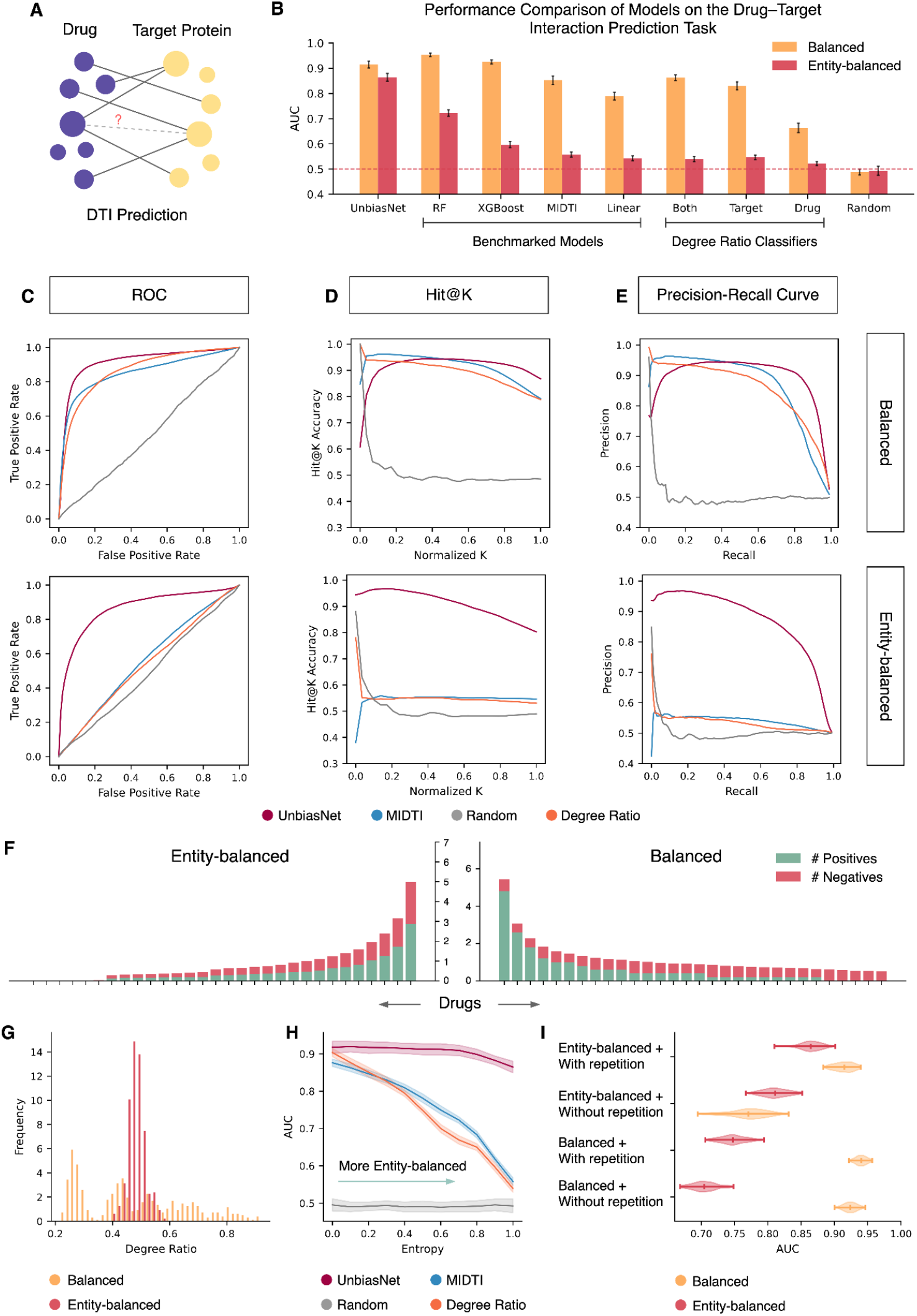
Evaluation of drug–target interactions prediction. **A**, Overview of the DTI prediction task. **B**, AUC of benchmarked models, baseline classifiers, and UnbiasNet under balanced and entity-balanced evaluation frameworks. While most models—including degree ratio baselines—perform well in the balanced framework, their performance drops sharply under entity-balanced evaluation. In contrast, UnbiasNet maintains strong performance across both frameworks. **C**, ROC curves for UnbiasNet, MIDTI, Random, and degree ratio models under the two evaluation schemes. **D**, Stratified Hit@K curves for the same models; the x-axis shows normalized K, and the y-axis shows Hit@K accuracy (see Methods). **E**, Precision–Recall curves for the same models. Results in **C**–**E** highlight that degree ratio bias cannot be detected by the choice of conventional evaluation metrics. **F**, Average number of positive and negative associations per drug across test datasets (ordered by total samples per drug); the x-axis shows a subset of drugs from the LuoDTI dataset, and the y-axis shows counts in balanced vs. entity-balanced frameworks. **G**, Histograms of drug degree ratios across balanced and entity-balanced test sets (zeros illustrated in another diagram, see Supplementary Fig. 4C). Together, **F** and **G** reveal that degree ratios are highly skewed in balanced datasets but corrected by entity-balanced sampling. **H**, AUC performance of models on test datasets with varying degrees of entity balance, quantified by entropy. As entropy increases, performance of all models declines except UnbiasNet, which remains stable. **I**, Ablation study of UnbiasNet, showing that both entity-balanced training sets and their diversity are critical for robust performance.

**Fig. 3.**
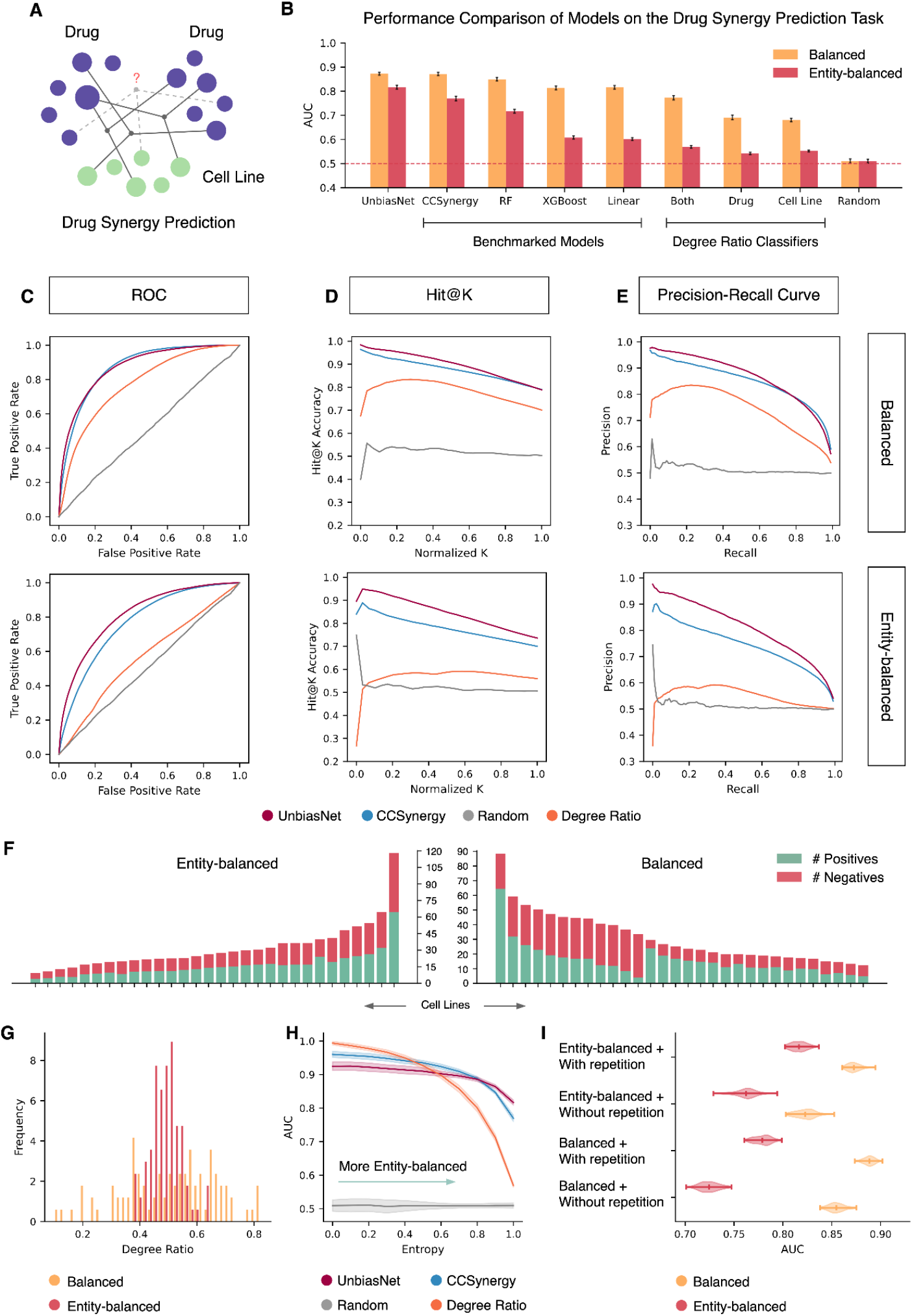
Evaluation of drug synergy prediction. **A**, Overview of the drug synergy prediction task. **B**, AUC of benchmarked models, baseline classifiers, and UnbiasNet under balanced and entity-balanced evaluation frameworks. Similar to the DTI setting, most models—including degree ratio baselines—perform well under the balanced framework but show substantial drops under entity-balanced evaluation. UnbiasNet maintains strong and consistent performance across both frameworks. **C**, ROC curves for UnbiasNet, CCSynergy, Random, and degree ratio models under the two evaluation schemes. **D**, Stratified Hit@K curves for the same models; the x-axis shows normalized K, and the y-axis shows Hit@K accuracy (see Methods). **E**, Precision–Recall curves for the same models. Results in **C**–**E** again illustrate that conventional metrics fail to expose degree ratio bias. **F**, Average number of positive and negative associations per cell line across test datasets (ordered by total samples per cell line); the x-axis shows a subset of cell lines, and the y-axis shows counts in balanced vs. entity-balanced frameworks. **G**, Histograms of degree ratios across cell lines in balanced and entity-balanced test sets. Together, **F** and **G** demonstrate that degree ratios are highly skewed in balanced datasets but corrected by entity-balanced sampling. **H**, AUC performance of models on test datasets with increasing entropy levels. **I**, Ablation study of UnbiasNet, showing that both the use of entity-balanced training sets and their diversity are essential for achieving robustness against shortcut learning.

### Performance Drop under the Entity-Balanced Evaluation Framework

For DTI prediction (Fig. 2A), we benchmarked MIDTI^18^ alongside conventional classifiers on the LuoDTI^27^ dataset. Since LuoDTI lacks negative labels, all non-positive associations were treated as negatives. Under conventional evaluation settings (Full Test and Balanced Evaluation), both classical and deep learning models achieved consistently high performance (Fig. 2B and Supplementary Fig. 1A). However, the strong performance of naïve baselines that rely solely on degree ratio information indicates that much of this apparent success may be driven by shortcut learning. Strikingly, even extremely simplistic baselines that relied exclusively on degree ratios from a single entity type—for instance, considering only target proteins—while being entirely blind to the complementary entity, such as drugs, achieved performance comparable to that of sophisticated state-of-the-art models.

This bias persists across different evaluation metrics, which fail to reveal the influence of degree ratios (Figs. 2C-E and Supplementary Figs. 1-2). Whether evaluated using ROC curves, Precision–Recall curves, or rank-based metrics such as Hit@K, the degree ratio classifier achieves performance comparable to more complex models. This demonstrates that the choice of evaluation metric alone is insufficient to reveal the presence of shortcut learning driven by degree ratio bias.

Under the Entity-Balanced Evaluation framework, however, all models—both benchmark and baseline—showed substantial performance drops, confirming that their earlier success was largely driven by polarity bias (Figs. 2B–E). Degree ratio baselines, which previously matched or outperformed advanced models, fell close to random once entity polarity was neutralized, exposing their complete reliance on shortcut features. MIDTI also showed a sharp decline, with its performance converging toward that of the baselines. This indicates that much of MIDTI’s apparent advantage under conventional settings stemmed from exploiting the same polarity bias, rather than capturing genuine biological patterns.

### Degree Ratio Bias Explains Performance Drop

Degree ratio analysis explains both the inflated performance of models under conventional evaluation frameworks and their performance decline under the entity-balanced evaluation framework (Figs. 2F–G and Supplementary Fig. 3). In datasets constructed using conventional protocols, many entities appear almost exclusively in either positive or negative associations, or else exhibit highly skewed degree ratios. Since this signal is shared between training and test sets, models readily learn to exploit it, producing deceptively high scores. For example, if a drug appears only in positive associations during training, the model may conclude that the drug is intrinsically linked to positive outcomes. Consequently, it predicts positive associations whenever the drug is encountered, regardless of the target—behavior consistent with our one-sided degree ratio baselines. Thus, because this spurious signal also persists in the test sets, models can achieve deceptively high yet fundamentally flawed performance.

By applying our entity-balance sampling algorithm, we constructed datasets in which the number of positive and negative associations for each entity is approximately equal (Figs. 2F–G and Supplementary Fig. 3). This design allows us to directly test whether a model relies on degree ratio shortcuts. Returning to the earlier example, if a model has learned to associate a specific drug exclusively with positive outcomes, its performance will collapse on an entity-balanced test set, since half of the associations involving that drug now belong to the opposite class. In this way, entity-balanced evaluation provides a clear diagnostic tool to distinguish genuine relational learning from shortcut-driven predictions.

### UnbiasNet Resists Degree Ratio Shortcut Learning

Our proposed model, UnbiasNet, achieved the strongest performance under the entity-balanced evaluation framework while simultaneously maintaining competitive results under conventional settings (Figs. 2B–E). Unlike other methods, the gap between its performance across the two frameworks was small, indicating limited reliance on degree ratio–based shortcut learning. To examine this robustness more systematically, we generated test sets with varying levels of entity balance, quantified by association entropy (see Methods). As the entropy increased—reflecting a more entity-balanced distribution of positive and negative associations—the performance of all competing models, including deep learning methods and classical classifiers, consistently declined (Figs. 2H and Supplementary Fig. 2B). By contrast, UnbiasNet maintained stable performance across these increasingly stringent conditions. Notably, the degree ratio classifier, which initially performed best under low-entropy conditions where polarity was strong, rapidly deteriorated as entity balance improved. In sharp contrast, UnbiasNet remained largely unaffected by the level of entropy, underscoring its ability to capture genuine relational signals rather than superficial statistical biases.

Our ablation study showed that both the use of entity-balanced training datasets and their diversity are critical for achieving robustness against shortcut learning (Fig. 2I). Relying on a single entity-balanced dataset sampled from the training pool does not provide sufficient variability for the model to effectively distinguish positive from negative associations, and in fact leads to reduced performance under conventional evaluation (Fig. 2I). Similarly, employing multiple non-entity-balanced datasets yields only marginal gains. In contrast, UnbiasNet integrates both entity balance and dataset diversity, resulting in substantially improved performance across both conventional and entity-balanced evaluation frameworks.

### Degree Ratio Shortcut Learning Extends Beyond Pairwise Entities

Building on our findings from drug–target interactions, we next turned to the more complex setting of drug synergy prediction, which involves three entities–two drugs and a cell line– as input (Fig. 3A). We benchmarked CCSynergy^16^ alongside conventional classifiers on the Sanger^28^ dataset. Unlike LuoDTI, this dataset includes experimentally validated negative associations, providing a more reliable basis for evaluation. The dataset was symmetrized with respect to the two drugs, ensuring that their positions within each triplet are interchangeable. Consequently, the degree ratio profiles of the first and second drugs are identical, simplifying the analysis of degree ratio effects in this higher-order association prediction task.

The findings mirrored those from the drug–target interaction task (Figs. 3B–I, Supplementary Figs. 4–6), though the performance decline under the entity-balanced evaluation framework was less pronounced than in the LuoDTI dataset (Figs. 3B–E, Supplementary Figs. 4–5). This milder drop reflects the more balanced distribution of entities in the Sanger dataset (Figs. 3F–G, Supplementary Fig. 6). Notably, CCSynergy displayed stronger resilience to degree ratio shortcut learning, a robustness partly attributable to the stratified testing strategy employed in its original paper^16^, which reduced dependence on degree ratios. In contrast, models such as XGBoost experienced sharp declines, underscoring that susceptibility to shortcut learning persists even when degree ratio signals are weaker. By contrast, UnbiasNet’s training strategy once again alleviated this reliance on unintended features (Figs. 3H–I). Together, the Entity-Balanced Evaluation framework and UnbiasNet demonstrated consistent performance in this setting, reinforcing their general applicability to multi-input association prediction problems.

## Conclusions

Standard evaluations serve as guidelines for the research community, shaping how new models and methods are developed to address open problems. However, when evaluation methodologies are flawed yet widely adopted, they risk steering progress in misleading directions. One key source of distortion arises from shortcut features hidden in curated biological datasets. These artifacts are dataset-specific and rarely generalize to real-world scenarios. Guided by the principle of least effort, models tend to exploit such shortcuts, since they are easier to learn than the intended biological features^21^. This makes it essential for the community to critically examine the datasets they use and remain vigilant about potential shortcut opportunities that could undermine fair evaluation.

Multi-input association prediction encompasses a wide spectrum of bioinformatics tasks but is often undermined by a pervasive shortcut signal: the degree ratio. To address this challenge, we present two complementary contributions. First, we propose an entity-balanced evaluation framework that mitigates shortcut learning by explicitly controlling for degree ratio bias, ensuring that model performance reflects genuine relational learning rather than superficial statistical patterns. Unlike conventional out-of-distribution splits, this framework remains practical for graph-based models, extends naturally to higher-order associations, and leverages the full set of negatives through entity-balanced subsampling. Second, we introduce UnbiasNet, a model-agnostic training strategy that constructs diverse entity-balanced sub-training sets, thereby systematically removing access to shortcut features. This approach demonstrates robustness to degree ratio bias while preserving strong performance under conventional evaluations. Together, these contributions establish a rigorous foundation for evaluating model limitations and promoting the development of methods that capture more meaningful biological signals. Beyond drug–target and drug synergy prediction, we expect these advances to generalize broadly across multi-input problems in computational biology.

## Methods

### Multi-input Association Prediction

We formalize biological association prediction as a hyperedge prediction problem within a multi-entity hypergraph. Let *V*_1_, *V*_2_, ⋯, *V*_*n*_ denote sets of entities belonging to different biological categories (e.g., drugs, proteins, cell lines). An association corresponds to a tuple (*v*_1_, *v*_2_, ⋯, *v*_*n*_ ) ∈ *V*_1_ × *V*_2_ × ⋯ × *V*_*n*_, which may be labeled as positive (associated) or negative (non-associated). We denote this as:

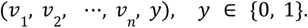

The collection of known associations and non-associations can be represented as a labeled hypergraph *G* = (*V, E*), where:

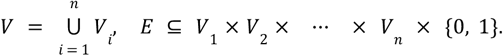

We refer to this labeled hypergraph as an Association Network, where each hyperedge represents an observed relationship among a group of entities.

Each entity *v* ∈ *V*_*i*_ is represented by a feature vector *x*_*v*_ ∈ *X*_*i*_, where *X*_*i*_ is the feature space of the i-th entity type. The objective is to learn a function:

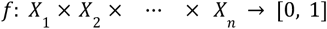

that estimates the probability of association for any tuple of entities based on their features. In drug–target interaction (DTI) prediction, a binary association links a drug from *V*_1_ with a protein from *V*_2_, and in drug synergy prediction, a ternary association involves two drugs from *V*_1_ = *V*_2_ and a cell line from *V*_3_ .

### The Entropy of an Association Network

To assess the polarity of entity associations within the graph, we introduce a measure of *association entropy*. Each entity may participate in both positive and negative associations, which are unevenly distributed across the network. For a node *v* ∈ *V*_*i*_, let *p*_*v*_ denote the number of positive associations (hyperedges labeled *y* = 1) in which *v* participates, and let *n*_*v*_ denote the number of negative associations (hyperedges labeled *y* = 0) involving *v*. The positive ratio is computed as:

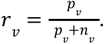

We then define the association entropy of node *v* as:

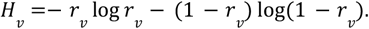

This entropy quantifies the uncertainty in the polarity of an entity’s associations, peaking at *r*_*v*_ ≈ 0. 5 and minimizing when associations are highly skewed toward either label. The aggregated entropy for entity set *V*_*i*_ ⊂ *V*, is given by the weighted average:

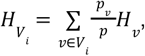

where *p* is the total number of positive associations. Finally, the entropy for the association network *A* is computed as the mean of the entropies across all entity types:

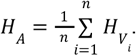

### Entity-Balanced Sampling Algorithm

To mitigate polarity imbalance and reduce the degree ratio bias, we developed the Entity-Balanced Sampling (EBS) algorithm. The method aims to ensure that each entity in the dataset appears in a roughly equal number of positive and negative associations. Given an association network *G* = (*V, E*) and hyperparameters for simulated annealing, EBS produces a subset 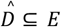 in which the per-entity label distribution is approximately balanced, such that the network entropy 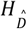 approaches 1. The procedure consists of two stages: an iterative negative sampling step guided by degree-ratio distributions, followed by refinement through simulated annealing.

#### Iterative Negative Sampling

This stage constructs an initial dataset 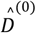 by augmenting the full set of positive associations *D*^+^ with a sampled subset of negatives 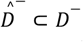. Here, *D*^−^ denotes the set of all possible negative associations. The goal is to sample |*D*^+^ | negatives such that the resulting polarity for each entity is more balanced. To guide the sampling process, we define a dynamic weight matrix *W*, which assigns scores to candidate negatives based on current entity-level imbalance. Specifically, for a candidate negative association *a* = (*v*_1_, *v*_2_, ⋯, *v*_*n*_, 0) ∈ *D*^−^, the weight is defined as:

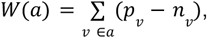

where *p*_*v*_ and *n*_*v*_ denote the number of positive and negative associations involving entity *v* currently observed in the combined set 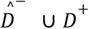. This scoring mechanism favors associations involving entities with skewed polarity (i.e., more positives than negatives), promoting balance as sampling progresses.

At each step, one negative is sampled from the remaining candidates in 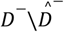 with probability proportional to *W*, added to 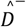. The weights are then recomputed to reflect the updated entity counts. This process is repeated until 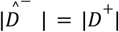, resulting in a more balanced initialization.

The output of this stage is the combined dataset:

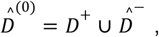

which serves as the starting point for the simulated annealing refinement. The Iterative Negative Sampling algorithm is presented in *Algorithm 1*.

##### Algorithm 1

Iterative Negative Sampling

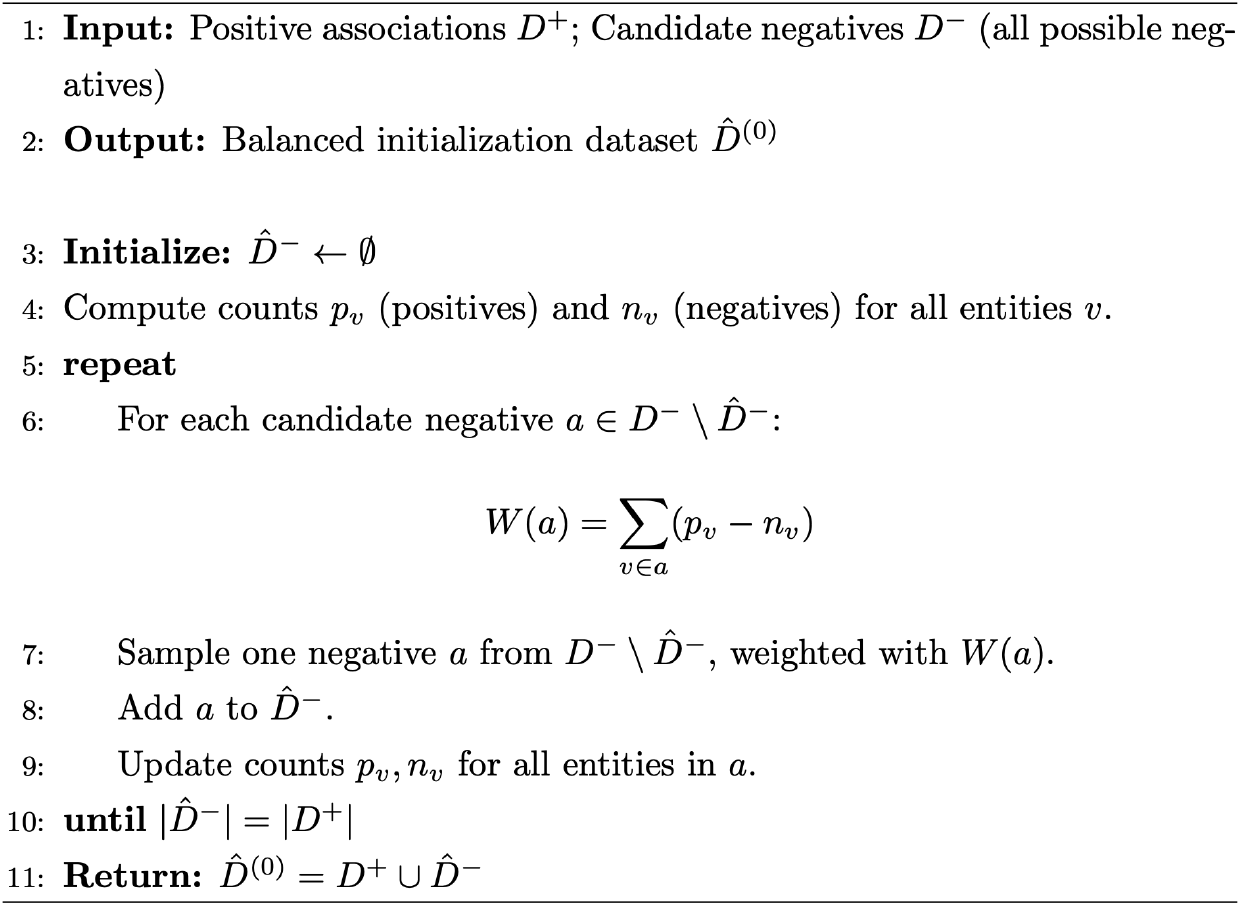

#### Simulated Annealing

We then refine 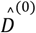 using simulated annealing. At each iteration *t*, we define the current dataset 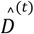 and evaluate an objective function that rewards polarity balance while discouraging excessive reduction in dataset size:

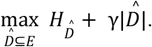

Here, 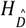 is the network-level association entropy as previously defined, and γ > 0 is a regularization coefficient that penalizes reductions in the number of samples, encouraging solutions that retain a sufficiently large and diverse dataset. At each iteration *t*, the algorithm selects one of two update moves: either removing one positive and one negative association from 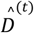, or adding one new positive and one new negative association from the candidate pool 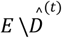. Moves that improve the objective are accepted, while occasional worse moves are allowed with decreasing probability controlled by a temperature schedule.

### Evaluation Frameworks

We examine three evaluation strategies: two widely adopted in the literature and one novel approach that we propose to explicitly reduce entity-level bias.

#### Full Test Evaluation

In this conventional approach, the dataset is randomly partitioned into training and test sets using a fixed split ratio^16^. Evaluation is performed on the complete test set, preserving the natural class imbalance observed in the data.

#### Balanced Evaluation

To reduce the impact of global class imbalance, this strategy constructs a balanced test subset from the fixed test set by uniformly sampling negative associations to match the number of positives^18^. Evaluation metrics are computed across multiple such subsets to reduce sampling variance. For example, the area under the ROC curve (AUC) is reported as an average:

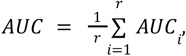

where *r* denotes the number of sampled sub-test sets, and *AUC*_*i*_ is the AUC score obtained from the *i*-th sub-test evaluation.

#### Entity-Balanced Evaluation

We propose Entity-Balanced Evaluation, a stricter framework that controls for polarity at the level of individual entities. Beginning with a fixed train–test split, we generate sub-test sets using the EBS algorithm, ensuring that each entity is represented by approximately equal numbers of positive and negative associations. Model performance is then reported as the average across multiple such sub-test sets.

### Stratified Hit@K

The Hit@K accuracy metric measures a model’s ability to prioritize true associations within its highest-ranked predictions. This metric is particularly valuable when negative samples may include uncertain or mislabeled cases. Formally, given a ranked list of predicted association scores, Hit@K accuracy is defined as the proportion of true positives that occur within the top *K* entries. By varying *K*, we obtain a Hit@K curve that provides a more comprehensive view of model performance. We employed this curve across all evaluation frameworks to compare their outcomes. Within each framework, multiple Hit@K curves are generated and subsequently aggregated as follows. For each sub-test set, we compute a Hit@K curve by ranking the predicted associations and calculating Hit@K scores for *K = 1* up to the total number of positives in that subset. Since the number of positives differs across sub-test sets, we normalize the x-axis of each curve to the interval [0, 1], representing relative ranking positions. All normalized curves are then interpolated over a common grid and averaged to yield the final Stratified Hit@K curve.

### Degree Ratio Classifiers

We introduce a class of baseline models known as Degree Ratio Classifiers. These classifiers rely exclusively on the polarity of individual entities—without considering any relational or contextual features—to predict the existence of an association.

#### Single-Entity Degree Ratio Classifier

A *V*_*i*_ -Degree Ratio Classifier focuses on a single entity type *V*_*i*_ and uses the degree ratio of an entity *v*_*i*_ ∈ *V*_*i*_ to score a given association:

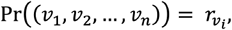

where 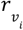 denotes the degree ratio of node *v*_*i*_ . This model assigns the same score to all associations that include a given *v*_*i*_, ignoring other entities in the tuple. For example, in a drug–target prediction task, a Drug-Degree Ratio Classifier assigns the same score to all pairs containing a given drug, based only on its degree ratio, entirely ignoring the specific target.

#### Weighted Mean Degree Ratio Classifier

To extend beyond single-entity scoring, we define the Weighted Mean Degree Ratio Classifier, which computes a weighted average of degree ratios across all entities in a tuple. For an association (*v*_1_, *v*_2_, ⋯, *v*_*n*_ ), the prediction is:

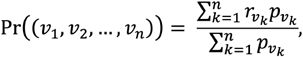

where *r*_*v*_ and *p*_*v*_ denote the degree ratio and the number of positive associations of node *v*. In this formulation, each entity’s contribution is weighted by its prevalence in positive associations.

### UnbiasNet Learning Framework

To mitigate model bias toward the degree ratio shortcut learning, we introduce an adjustment to the training process that leverages our EBS algorithm. Rather than relying on the naturally imbalanced dataset or a single balanced version, which risks omitting important negative or positive examples, we propose a dynamic training regime based on a collection of entity-balanced sub-training sets. These subsets are constructed using the EBS algorithm, each initialized with a different random seed to enhance diversity. Let *k* denote the number of entity-balanced sub-training sets generated. During training, the model cycles through these subsets: at epoch *i*, the training set used is the (*i mod k*)-th balanced subset.

### Models

We evaluated two state-of-the-art models: CCSynergy^16^ for drug synergy prediction and MIDTI^18^ for drug–target interactions. Their feature representations were also used with standard classifiers to separate the effects of features from model architecture.

#### CCSynergy

CCSynergy^16^ is a deep learning-based framework designed for context-aware prediction of anti-cancer drug synergy. It leverages bioactivity signatures from the Chemical Checker (CC), which captures drug characteristics across multiple biological levels. In our study, we specifically used biological processes features from the CC catalog to represent each drug. These features integrate network-level and pathway-level information, offering a functionally relevant view of compound activity. For modeling the cellular context, we employed transcription factor (TF) activity profiles as features for cell lines.

#### MIDTI

MIDTI^18^ is a state-of-the-art model for drug–target interaction prediction based on a multi-view similarity network fusion and a deep interactive attention mechanism. MIDTI constructs heterogeneous networks capturing various biological relationships and fuses multiple similarity views for drugs and targets using an attention-enhanced fusion strategy. Embeddings are then learned using Graph Convolutional Networks (GCNs) over several network types: drug similarity, target similarity, drug–target bipartite, and drug–target heterogeneous networks. Final association scores are computed via a multilayer perceptron applied to the concatenated embeddings of each drug–target pair.

During reimplementation, we identified a critical flaw in the original MIDTI design: drug and target features were derived from the complete association matrix prior to train–test splitting, resulting in data leakage. To address this, we developed a corrected variant, which applies feature extraction strictly within the training partition, ensuring leakage-free evaluation.

### Datasets

We evaluated model performance on two widely used benchmark datasets, each representing a distinct class of association prediction problems: drug synergy prediction (three entities) and drug–target interaction (two entities).

#### Sanger Drug Synergy Dataset

For the drug synergy prediction task, we used a curated subset of the large-scale screening dataset published by the Sanger^28^ Institute, as processed in the CCSynergy^16^ study. The original dataset includes 2,025 drug combinations tested across 125 cancer cell lines, totaling over 108,000 triplets. Following the filtering criteria introduced in CCSynergy—specifically the availability of Chemical Checker (CC) signatures for drugs and comprehensive cell line features—we used the refined subset consisting of 62 drugs, 1,177 drug pairs, and 93 cell lines, yielding a total of 46,748 distinct triplets^16^.

Each triplet was assigned a binary label (synergistic or non-synergistic) based on changes in potency and efficacy relative to Bliss independence expectations. A triplet was considered synergistic if at least half of its replicates showed synergy^16^. Consistent with prior work, we treated drug order as symmetric, duplicating each triplet with the drug positions swapped to enforce order invariance during training.

#### LuoDTI Dataset

For the drug–target interaction task, we used the dataset introduced by Luo et al.^27^ This dataset includes 1,923 known drug–target interactions among 708 drugs and 1,512 protein targets. Since it only contains positive examples, we followed standard practice by treating all unknown drug–target pairs as candidate negatives, resulting in a large number of potential negative instances.

## Data Availability

The datasets used in this study were obtained from previously published resources. Specifically, the LuoDTI dataset and its associated features for the MIDTI model were obtained from https://github.com/XuLew/MIDTI, and the Sanger dataset together with the features for the CCSynergy model were obtained from https://github.com/RzgarHosseini/CCSynergy. All data generated and used in this work are available at https://doi.org/10.6084/m9.figshare.30119653.v1.

## Code Availability

All code used for data processing, analysis, and evaluation, as well as tutorials for reproducing the results, are available at: https://github.com/sobhanAhmadian/netbalance.

