## Supplemental Figures 1-6 for "Reliable Evaluation and Learning in Multi-input Biological Association Prediction"

### Supplementary Materials for "Reliable Evaluation and Learning in Multi-input Biological Association Prediction"

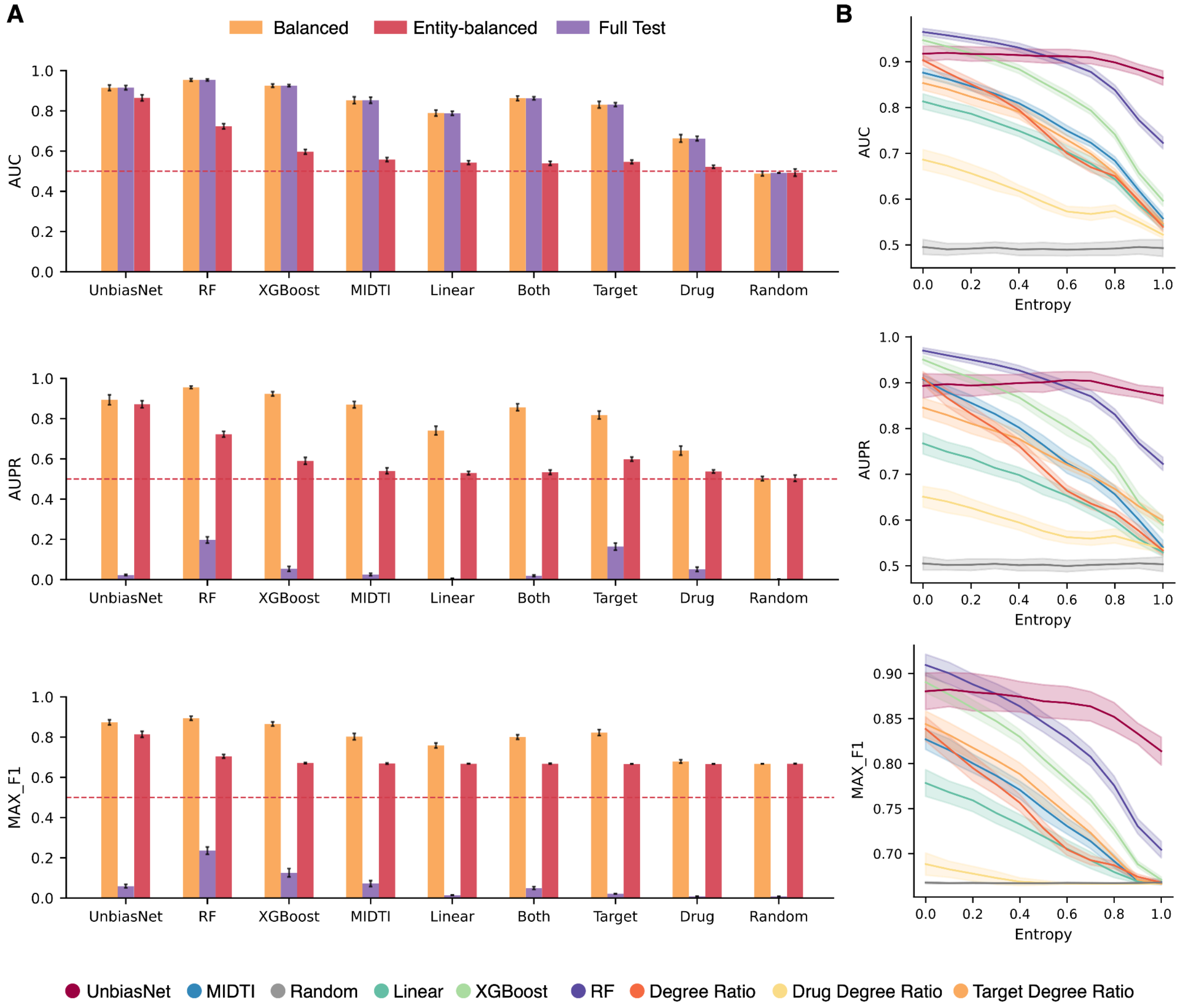


Sup. Fig. 1 | **Additional evaluation of drug–target interaction prediction. A**, AUC, AUPR, and Max F1 scores of benchmarked models, baseline classifiers, and UnbiasNet under balanced, full-test, and entity-balanced evaluation frameworks. AUPR and Max F1 follow the same trend as AUC in balanced and entity-balanced settings. In contrast, most models show near-zero AUPR and Max F1 under the full-test framework, where negatives vastly outnumber positives. Because many negatives in full-test datasets are uncertain, results from this framework are less reliable. **B**, AUC, AUPR, and Max F1 scores for the same models evaluated on test datasets with varying levels of entity balance. As entropy increases, performance of all models declines, whereas UnbiasNet remains comparatively stable.

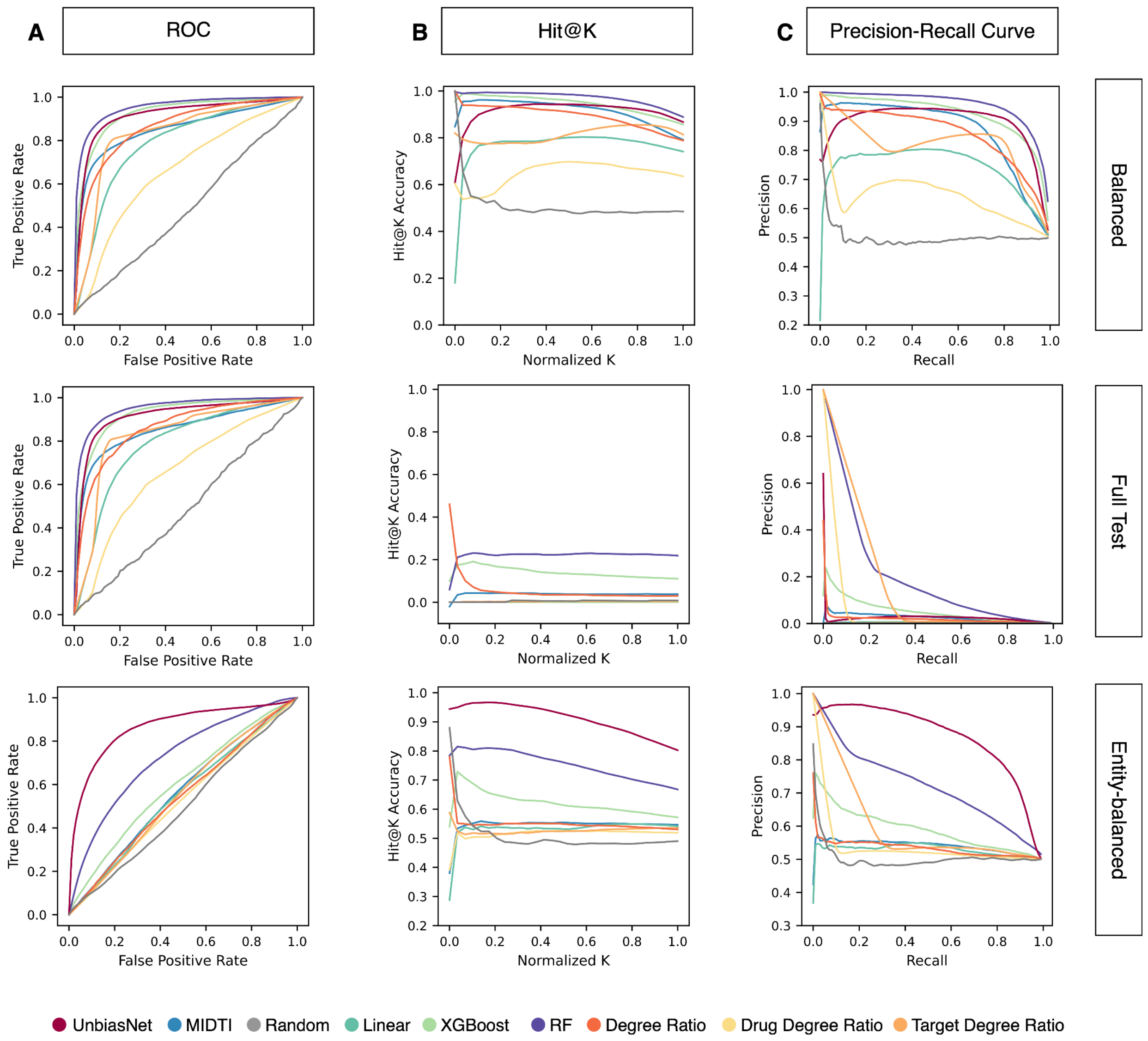


Sup. Fig. 2 | **Additional evaluation of drug–target interaction prediction. A**, ROC curves for benchmarked models, baseline classifiers, and UnbiasNet under the three evaluation schemes. **B**, Stratified Hit@K curves for the same models; the x-axis shows normalized K, and the y-axis shows Hit@K accuracy. **C**, Precision–Recall curves for the same models.

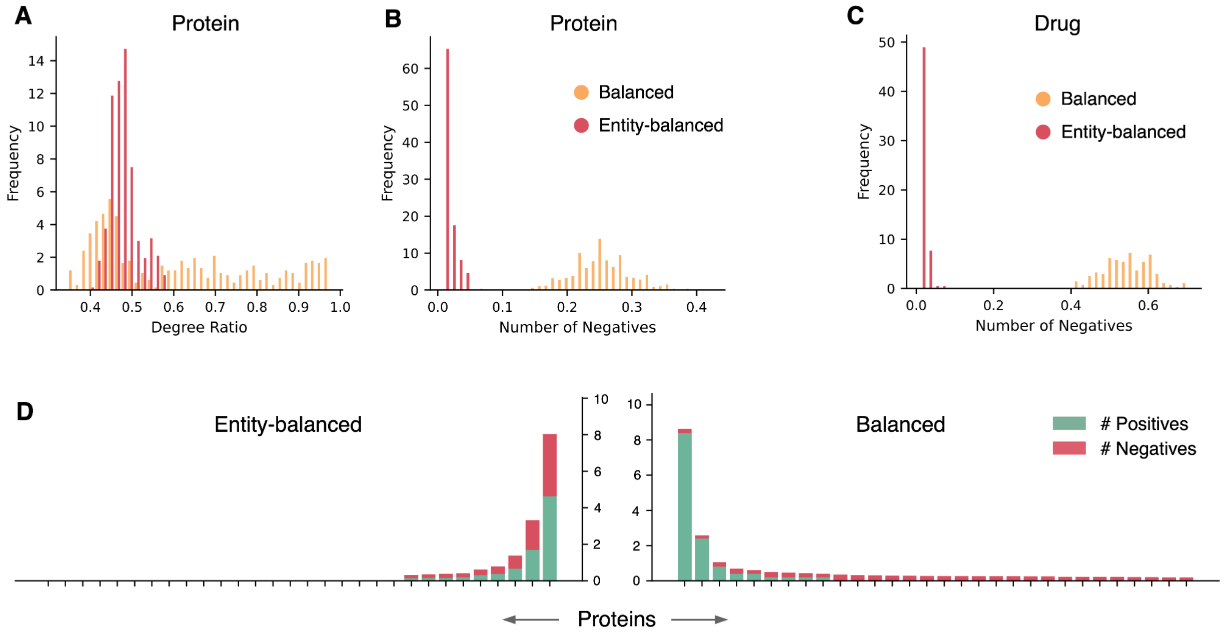


Sup. Fig. 3 | **Additional degree ratio analysis of drug-target interaction prediction. A**, Histograms of protein degree ratios in balanced and entity-balanced test sets (excluding zeros). **B**, Histogram of the average number of negative associations for proteins with zero positive associations; values are closer to zero in entity-balanced datasets. **C**, Same as **B**, but for drugs. **D**, Average number of positive and negative associations per protein across test datasets, ordered by total samples per protein; the x-axis shows a subset of proteins from the LuoDTI dataset, and the y-axis shows counts in balanced vs. entity-balanced frameworks.

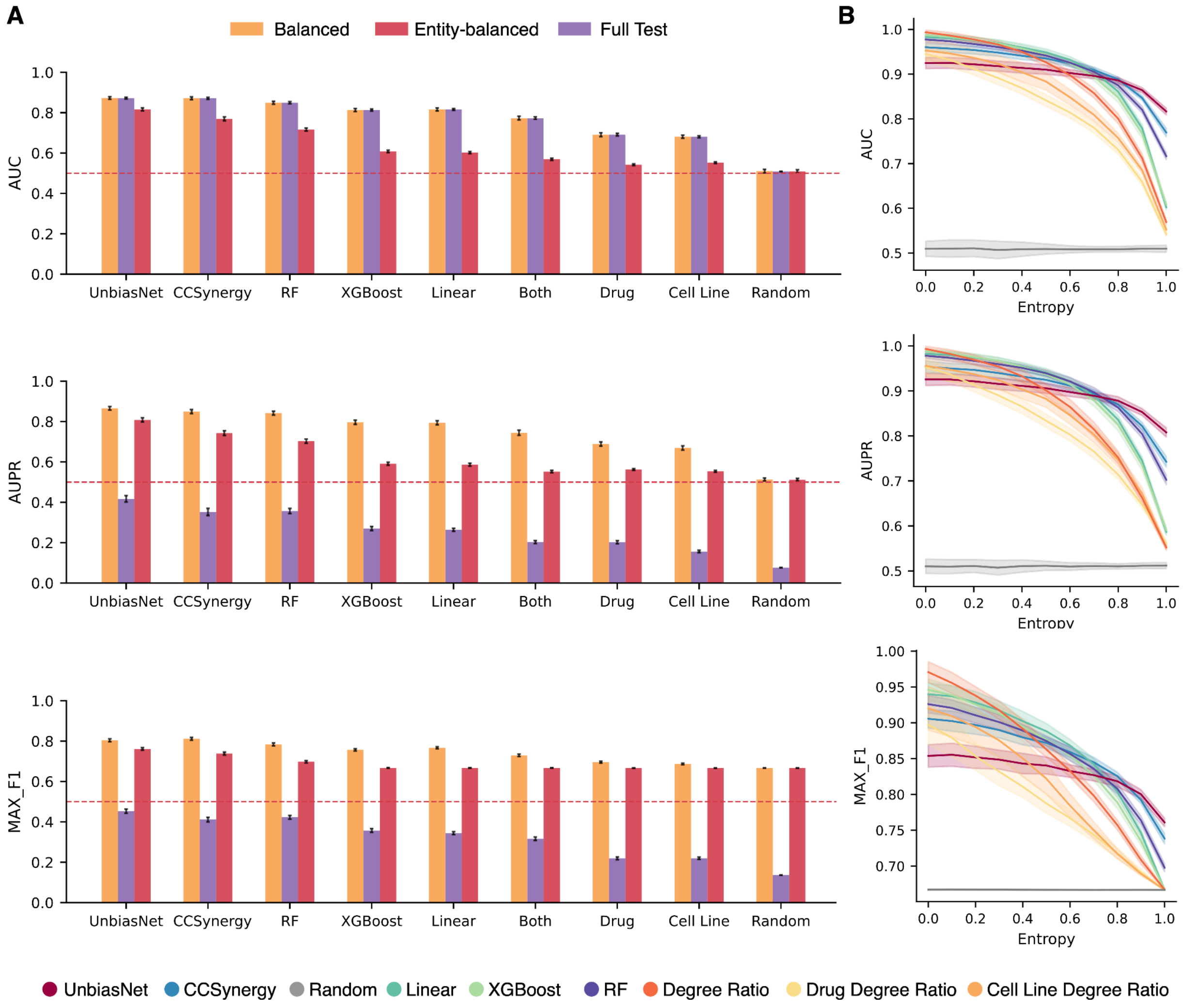


Sup. Fig. 4 | **Additional evaluation of drug synergy prediction. A**, AUC, AUPR, and Max F1 scores of benchmarked models, baseline classifiers, and UnbiasNet under balanced, full-test, and entity-balanced evaluation frameworks. As in the DTI task, AUPR and Max F1 track AUC in balanced and entity-balanced settings, where positives and negatives are matched. In the full-test framework—where negatives greatly outnumber positives—models show reduced AUPR and Max F1, but the relative ranking of methods remains consistent with the other evaluation settings. Unlike LuoDTI, the Sanger dataset contains true negative samples, making full-test results more reliable. **B**, AUC, AUPR, and Max F1 scores for the same models across test datasets with varying levels of entity balance. As entropy increases, performance declines for all models except UnbiasNet, which remains comparatively stable.

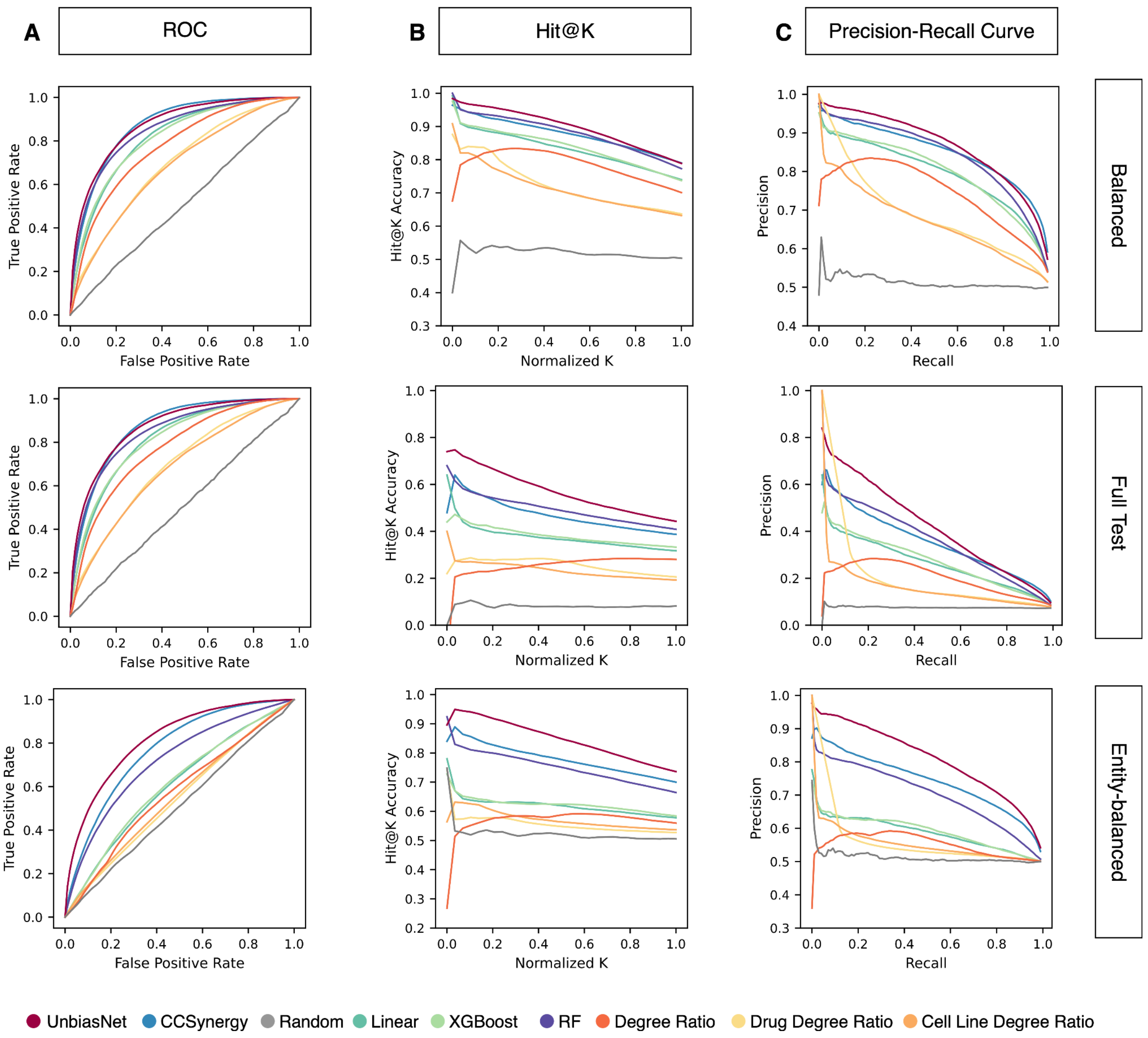


Sup. Fig. 5 | **Additional evaluation of drug synergy prediction. A**, ROC curves for benchmarked models, baseline classifiers, and UnbiasNet under the three evaluation schemes. **B**, Stratified Hit@K curves for the same models; the x-axis shows normalized K, and the y-axis shows Hit@K accuracy. **C**, Precision–Recall curves for the same models.

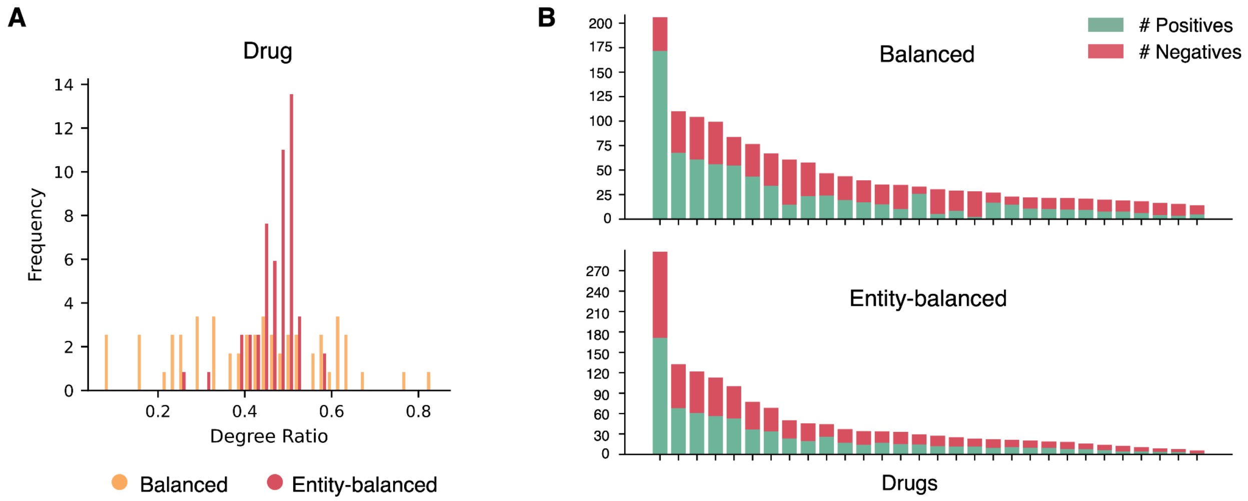


Sup. Fig. 6 | **Additional degree ratio analysis of drug synergy prediction. A**, Histograms of drug degree ratios in balanced and entity-balanced test sets. **B**, Average number of positive and negative associations per drug across test datasets, ordered by total samples per drug; the x-axis shows a subset of drugs from the Sanger dataset, and the y-axis shows counts in balanced vs. entity-balanced frameworks.
